# Saliva sampling is an excellent option to increase the number of SARS CoV2 diagnostic tests in settings with supply shortages

**DOI:** 10.1101/2020.06.24.170324

**Authors:** Joaquín Moreno-Contreras, Marco A. Espinoza, Carlos Sandoval-Jaime, Marco A. Cantú-Cuevas, Héctor Barón-Olivares, Oscar D. Ortiz-Orozco, Asunción V. Muñoz-Rangel, Manuel Hernández-de la Cruz, César M. Eroza-Osorio, Carlos F. Arias, Susana López

## Abstract

As part of any plan to lift or ease the confinement restrictions that are in place in many different countries, there is an urgent need to increase the capacity of laboratory testing for SARS CoV-2. Detection of the viral genome through RT-qPCR is the golden standard for this test, however, the high demand of the materials and reagents needed to sample individuals, purify the viral RNA, and perform the RT-qPCR test has resulted in a worldwide shortage of several of these supplies. Here, we show that directly lysed saliva samples can serve as a suitable source for viral RNA detection that is cheaper and can be as efficient as the classical protocol that involves column purification of the viral RNA. In addition, it surpasses the need for swab sampling, decreases the risk of the healthcare personnel involved in this process, and accelerates the diagnostic procedure.

## INTRODUCTION

With the worldwide COVID-19 health emergency, there is an urgent need for rapid and reliable methods of diagnostic for SARS-CoV-2. The accepted golden standard for detection of this virus is the amplification of regions of the viral genome by RT-qPCR in nasopharyngeal and oropharyngeal swabs [1][2]. Unfortunately, given the enormous demand of the reagents needed to collect the biological samples, and to purify the viral RNA, there have been shortages of many of the reagents needed for the diagnostic tests. Swabs, viral transport media, and kits for viral RNA extraction are amongst the most common consumables that have become scarce, compromising the number of tests that can be done in many parts of the world.

Recently, several reports have demonstrated the possibility of using saliva instead of oral and nasal swabs to detect the genome of SARS-COV-2 [3][4][5]. Saliva collection also has many collateral benefits, including self-collection, what decreases the risk of healthcare workers in charge of taking the swabs, and does not require the use of PPE (personal protecting equipment) that has also become a scarce item in this pandemia [6][7]. In addition, the methods to extract the RNA from biological samples require the use of purification kits whose availability has also become limited due to the heavy worldwide demand.

In this report, we compared the RT-qPCR results from 253 paired samples obtained from saliva and swabs of ambulatory patients; the RNA in the swab samples was extracted using a commercial RNA purification kit, and the saliva samples were directly mixed with a lysis buffer, boiled, and used for the RT-qPCR protocol. We found a very good correlation of results between both types of samples, and propose that this method, which simplifies the sampling of patients, and accelerates the preparation of the RNA for the RT-qPCR test represents an excellent alternative that will facilitate to sample and diagnose a larger number of persons at a reduced cost.

## MATERIALS AND METHODS

### Sample collection

253 paired samples from oropharyngeal and/or nasopharyngeal swabs, and saliva were collected during a span of 30 days (from May 2^nd^ to 31^st^) by healthcare workers from the epidemiology department of the health ministry of the state of Morelos (Secretaría de Salud Morelos, SSM). All but 3 samples, were from ambulatory patients, the 3 exceptions were collected from hospitalized patients.

### Swab sampling

Oropharyngeal and nasopharyngeal swabs were taken from 71 patients, while a single oropharyngeal swab was taken from 182 patients. After their collection, swabs were placed in 2.5 ml of viral transport medium.

### Saliva Collection

Saliva was self-collected by patients that were asked to spit on several occasions into sterile urine cup containers until completing roughly 2-3 ml of saliva. No viral transport media, nor stabilizing agents, were added to the saliva samples.

After collection, both swab and saliva samples were stored and kept at 4°C until transported to the Institute of Biotechnology/UNAM for their analysis, which was within 24-36 hours after sample collection.

### Nucleic acid extraction and SARS-CoV-2 detection by RT-qPCR

Total RNA was extracted from swab samples using the QIAamp viral RNA mini kit (QIAGEN) following the manufacturerμs protocol, using 140 μl of viral transport medium from each swab, and the purified RNA was eluted in 60 μl of elution buffer.

Saliva samples were treated with the Quick Extract^TM^ DNA Extraction Solution (QE, Lucigen) by mixing 50 μl of saliva with 50 μl of the QE reagent and heating for 5 minutes at 95°C, the mixture was then cooled on ice and kept at 4°C until their use (within 1 hour of QE treatment). In saliva samples that had high viscosity, 1 volume of sterile phosphate-buffered saline (PBS) was added and mixed by repeated pippeting, and the diluted saliva sample was the extracted as mentioned above.

SARS-CoV-2 detection was performed using the Berlin protocol, using the reported oligos and probes for viral gene E and for human RNase P [19]. The RT-qPCR reactions were performed using the StarQ One-Step RT-qPCR (Genes 2 Life) kit, using 5 μl of the column extracted total RNA in a 20 μl of reaction mix, or 2.5 μl of the QE treated saliva into 22.5 μl of reaction mix. Samples were analyzed in an ABI Prism 7500 Sequence Detector System (Applied Biosystems) with the following thermal protocol: 50°C for 15 min, 95°C for 2 min and then 45 cycles of 95°C for 15 s and 60°C for 30 s. All samples with a Ct value equal or less than 38 were classified as positive.

### Determination of viral copy number

To determine the viral copy number in a sample, a standard curve was generated using a 10-fold serial dilution of an *in vitro* T7 RNA transcript that encodes the sequence recognized by oligonucleotides and probe for gene E. Briefly, the logarithm of concentration of each dilution was plotted against the Ct and the viral copy number from unknown samples was determined by extrapolating the Ct value onto the corresponding standard curve.

### Statistical analysis

Statistical analysis was performed using GraphPad Prism 6.0 (GraphPad Software Inc.) as described in the results section.

## RESULTS

### Detection of SARS-CoV-2 in paired swab and saliva samples

To evaluate if saliva is a good source of viral RNA for the RT-qPCR tests we determined the presence of the SARS-CoV-2 genome in paired saliva and swab samples from 253 ambulatory patients. All patients had two or more symptoms related to COVID-19 [8][9], 115 (45.4%) were male and 137 (54.1%) female, with a median age of 41 (+/−14.4) years. Samples were taken from ambulatory patients in the respiratory triage of the Tlaltenango health center, in Cuernavaca, Morelos. The RT-qPCR Berlin protocol was used to detect SARS-CoV-2, using only the primers and probe for gene E, since previous studies have shown a weak detection of viral RNA when the RdRp gene is probed [10][11]. As an internal control of RNA content in the samples, the RNase P gene was detected. Total RNA was purified from swabs using the QIAamp viral RNA mini kit; the RNA in saliva was directly obtained using the QE lysis buffer (Lucigen) and boiling for 5 min, as reported [12].

During the course of the study, and due to the shortage of swabs, the health center shifted temporarily from collecting two swabs per person (nasopharyngeal swab-NPS-plus oropharyngeal swab -OPS) to only one swab (OPS) per individual. From the 253 patients included in this study, two swabs were used in 71 (28%) of the cases, while a single OPS was taken from the other 182 (72%); irrespectively of the number of swabs collected, saliva samples were taken from all patients.

Of the 182 patients with a single swab collected, 80 (43.9%) were positive for SARS CoV2 either in the swab or saliva samples. Of these, 41 (51.2%) were positive in both types of samples, while 28 (35%) were only detected in saliva and not in the swab sample, and 11 (13.7%) were only positive in the OPS. In total, out of the 80 individuals found to be positive for the virus, 69 (86.2%) were correctly detected using saliva, while only 52 (65%) were detected with the OPS. (Table 1, Fig. 1).

**Table 1.**
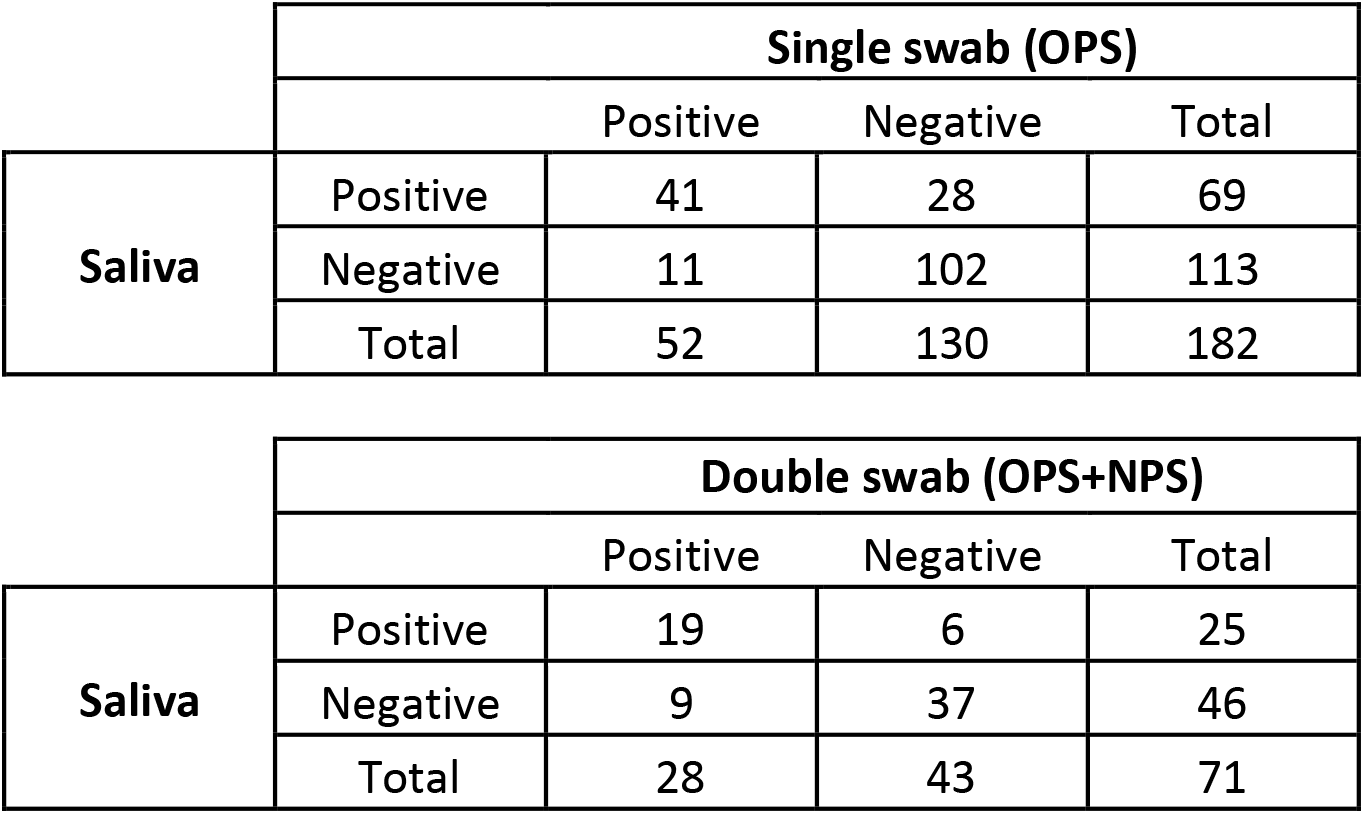
Summary of results obtained from parallel testing of swab and saliva samples from patients suspected of COVID-19.

|  | Single swab (OPS) |  |  |  |
| --- | --- | --- | --- | --- |
|  |  | Positive | Negative | Total |
| Saliva | Positive | 41 | 28 | 69 |
|  | Negative | 11 | 102 | 113 |
|  | Total | 52 | 130 | 182 |

|  | Double swab (OPS+NPS) |  |  |  |
| --- | --- | --- | --- | --- |
|  |  | Positive | Negative | Total |
| Saliva | Positive | 19 | 6 | 25 |
|  | Negative | 9 | 37 | 46 |
|  | Total | 28 | 43 | 71 |

**Figure 1.**
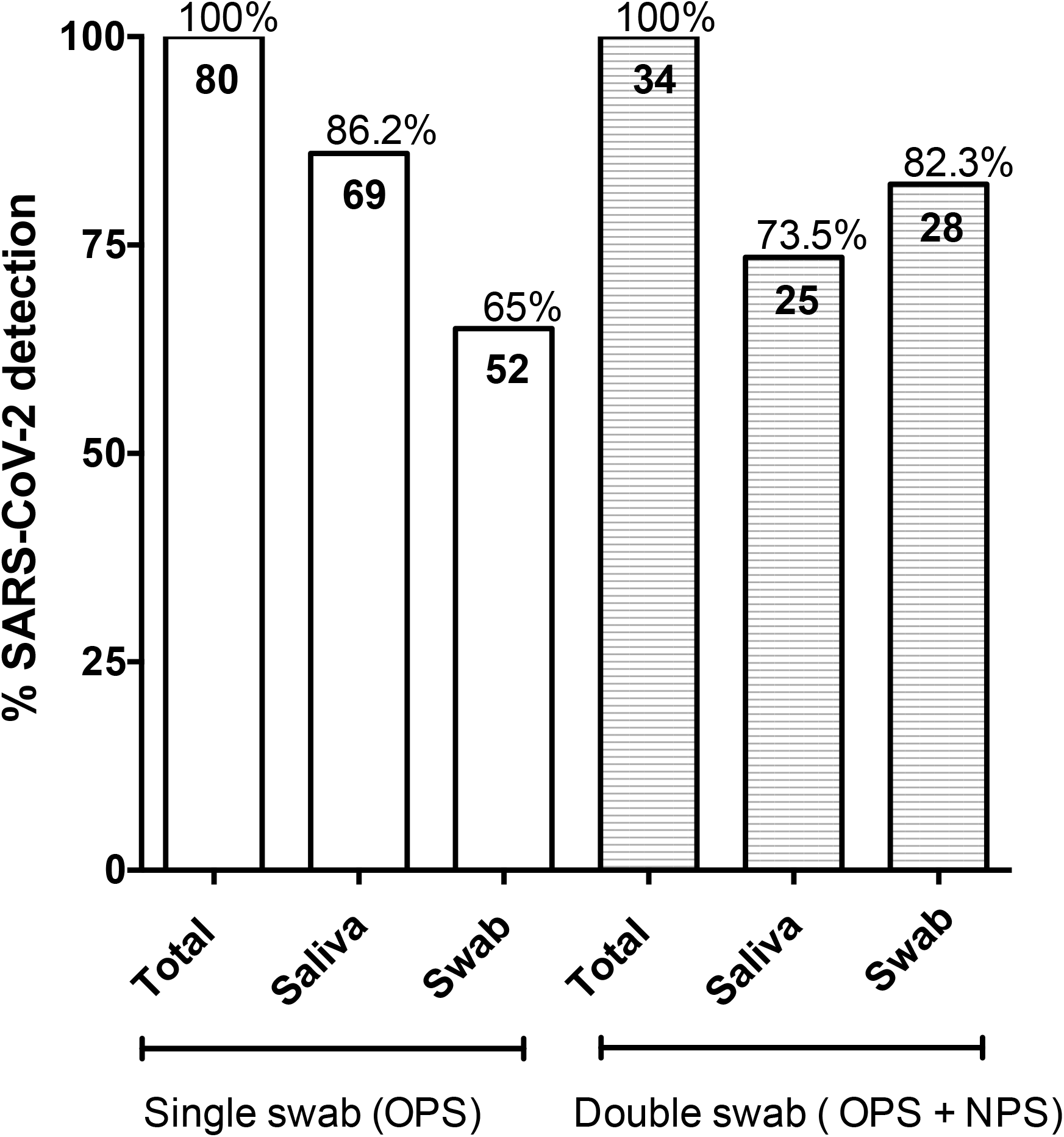
Detection of SARS-CoV-2 in paired swab and saliva samples. Percent number of positive samples detected in single OPS and saliva, or double (OPS + NPS) and saliva, as indicated. Data are extracted from Table 1.

On the other hand, 34 (47.8%) of the 71 patients with two swabs collected were found positive for SARS-CoV-2 in either the swabs or the saliva samples. Of these, 19 (55.8%) were positive both in swabs and saliva, while 6 (17.6%) were only positive in saliva, and 9 (26.4%) were only positive in the two-swab sample. In total, in this group of patients, of the 34 individuals detected as positive for the virus, 25 (73.5%) were identified by testing saliva, while 28 (82.3%) were positive by testing the swabs (Table 1 and Fig. 1).

### Quantitation of viral RNA

When the number of viral genome copies in the single OPS and saliva samples were compared, a significant difference in the geometric mean was detected, with saliva samples having a 1.9 log_10_ higher titer than that observed in the swabs (p<0.0024, Fig. 2A). This can be better appreciated when the viral copy number in paired swabs and saliva from the same patient, is plotted and represented as connecting lines (Fig. 2B); in 31 of the paired samples the number of viral copies was higher in saliva samples than in swabs. Human RNase P was used as an internal control of sampling quality; of interest, the comparison between the mean of Ct values obtained from OPS and saliva samples showed a difference of at least 6.8 Ct's between both types of samples (Fig. 2C), indicating that there is more cellular material in saliva, as reported in other studies[13]. The viral genome copy number in the double-swab and saliva samples was not statistically different, although a larger set of data would be needed to confirm these results (data not shown).

**Figure 2.**
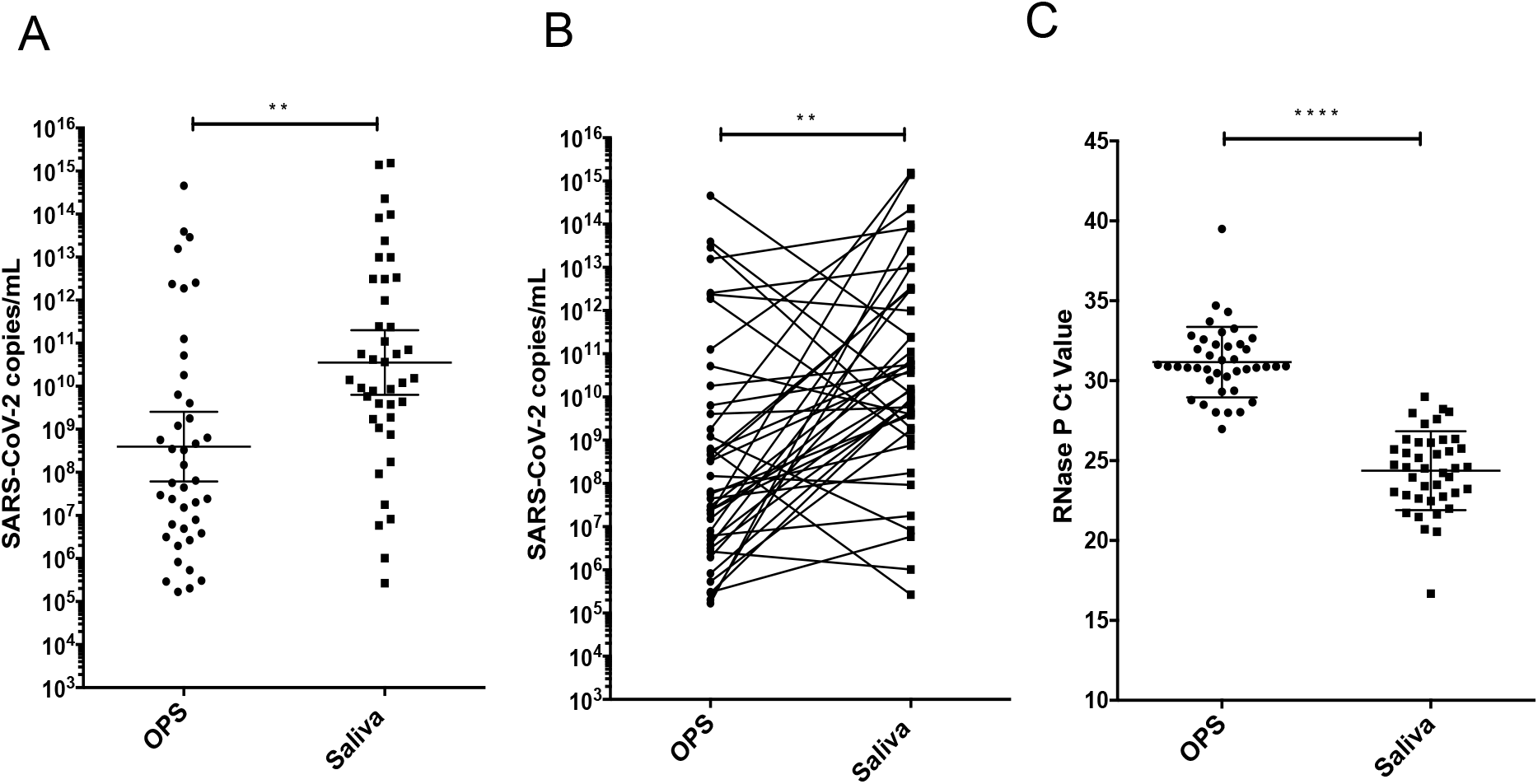
A high SARS-CoV-2 genome copy number is detected in saliva samples. A) Viral titer (viral copies/ml) detected in paired OPS and saliva samples B) Viral titer detected in paired OPS and saliva samples were represented by lines connecting both samples. Data were compared by a Wilcoxon test (p<0.0024); C) RT-PCR cycle threshold Ct values for RNase P detected in OPS and saliva samples. Data were compared by Wilcoxon test (p<0.0001). Bars represent the geometric median and 95% CI.

Taken together, these results suggest that that saliva is a good source for SARS-CoV-2 detection, especially when compared with a single OPS. Furthermore, it can be implemented for diagnostic tests using a simple QE buffer-based sample preparation in place of the column-based RNA purification method that is currently employed for swab analyses.

## DISCUSSION

In this study we analyzed 253 paired samples from OPS, or OPS and NPS and saliva. RNA purified from swabs using commercial column kits was compared with saliva samples directly lysed with QE buffer (surpassing the RNA extraction protocol), as source for the RT-qPCR assay. Although the coincidence rate between the single OPS and saliva samples was relatively low (51.2%), the saliva samples were clearly more efficient in detecting the virus when compared to single OPS samples (86.2% vs 65%). On the other hand, the efficiency of detection of the virus in saliva when compared to the double OPS and NPS was slightly lower (73.5% vs 82.3%), with a coincidence rate of 55.8%.

The reason for the low coincidence in the positive results obtained with swab and saliva samples is not clear. The failure of identification of SARS-CoV-2 in swabs, when the saliva samples were positive for the virus, could be due to bad swab sampling, what can be corroborated by the higher Ct values of RNase P detected in these samples (Fig. 2C), with the consequent low viral copy number. This is a major concern, since the medical personnel in charge of taking the samples frequently do not do it correctly for the risk associated with this process. It has been reported that oropharyngeal swabs have a lower viral titer compared with nasopharyngeal swabs [2]; thus, this could contribute to the discrepancies observed. Furthermore, it has also been previously reported that nasopharyngeal swabs have a lower viral titer than saliva samples [13], what can also contribute to explain our findings. On the other hand, the false negatives in saliva could be due to undetected problems during the collection, transport and or storage of the sample before its arrival to the laboratory.

SARS-CoV-2 has been detected in saliva at higher titers during the first days after the onset of symptoms, with the viral titer declining over time. It is not clear how long after the symptom onset the viral RNA can be detected in saliva, although some reports suggest a short period of detection (~13 days) as compared with nasopharyngeal swabs (~19 days) [14]. However, other reports have recently demonstrated the detection of viral RNA in saliva for longer periods of time (~20 days or longer) [3][15]. The patients included in this study were ambulatory, and according to their clinical interview had between 1 and 7 days (median of 4 days) of the onset of symptoms. We did not find a significative difference between the onset of symptoms and the results obtained from samples that were only positive in saliva versus those that were only detected in swabs.

Direct lysis of nasopharyngeal or oropharyngeal swab samples in viral transport medium using the QE buffer has been reported as a suitable method for direct RT-qPCR for SARS-COV-2 detection, with rates similar to methods based on column purification [16][12]. However, in our experience we have found a great variability in the results obtained using the QE lysis protocol when applied to swab samples, most likely due to variations in the material of the swabs used and to variations in the preparation of the viral transport medium employed (data not shown). In this regard, it has recently been reported that the composition of viral transport media can affect the detection of viral RNA from SARS-CoV-2 and other viruses [17] and, due to the scarcity of it, several laboratories have started to prepare their own transport media introducing an additional confusion factor. A similar situation occurs with the swabs, since in view of the scarce suitable materials, other materials are being employed, despite the fact that some of them are known to inhibit the RT-PCR reactions [18].

The use of saliva samples offers the advantage that no additives or transport media need to be used for their preservation or analysis if stored in cold and analyzed up to 36 h after their collection. Our results indicate that a rapid processing of saliva using direct lysis with QE buffer offers an excellent alternative to the current swab analysis that uses RNA column purification, since it is a sensitive, fast and cheap method that can be used for massive screening, in particular in those settings where common supplies needed for the classical methods are in shortage.

## ACKNOWLEDGMENTS

We are grateful to the healthcare workers of Servicios Estatales de Salud de Morelos for their invaluable help in collecting the samples, and to the personnel of the Laboratorio Estatal de Salud Pública del Estado de Morelos, for their support in the preparation and transporting of the samples. Part of the reagents used in this study were provided by the Instituto Nacional de Diagnóstico y Referencia Epidemiológica supported by INSABI. This work was supported by grant 314343 from CONACyT.

## Notes

### Competing Interest Statement

The authors have declared no competing interest.

